# Local nesting resources rather than floral food alter ground-nesting bee communities in grazed habitats

**DOI:** 10.1101/2024.08.19.608671

**Authors:** Shannon M. Collins, Laura A. Taylor, Karen W. Wright, Elinor M. Lichtenberg

## Abstract

1. Understanding how the availability of both trophic and non-trophic resources impacts key ecosystem service providers is crucial for informing conservation efforts. Non-trophic resources such as shelter can play critical roles in ecosystem function and resilience by affecting predation and pathogen rates, reproductive success, and interspecific interactions. However, non-trophic resources are often overlooked. This holds true for insect pollinators; impacts of shelter on community dynamics and pollination remain poorly understood.
2. Because the majority of bee species spend most of their lives underground, we explored whether nesting habitat characteristics are key drivers of ground-nesting bee abundance, diversity, and community composition. We additionally compared impacts of aboveground versus belowground nesting habitat characteristics, as the latter have received significantly less attention. Nesting habitat characteristics we considered included bare ground cover; and soil texture, compaction, and depth. We also included floral abundance and richness. We focused on rangelands, natural or semi-natural grasslands that are managed for livestock grazing, where management can dramatically alter trophic and non-trophic resources. Thus, we also investigated impacts of grazing management on bee habitat.
3. We found that belowground, but not aboveground, nesting habitat characteristics were important for ground-nesting bee assemblages. Ground-nesting bee abundance and richness were highest at sites with sandier and less compacted soils.
4. Our results suggest that livestock grazing management may indirectly impact ground-nesting bee abundance through impacts on floral abundance. Sites with longer grazing rest periods had higher floral abundance, and sites with higher floral abundance supported more ground-nesting bees (but not more bee species).
5. *Policy implications:* Our study highlights the importance of considering non-trophic resources in conservation and restoration management. Managers should be cautious in using an aboveground measure such as bare ground as an indicator of bee habitat quality. Rather, we recommend that belowground habitat characteristics such as soil texture and compaction be included in habitat quality assessments. Evidence-based rangeland conservation and restoration planning may need to manage for ground-nesting bees’ preferred levels of soil compaction. Additionally, management-insensitive habitat characteristics such as soil texture may be important considerations for site prioritization in pollinator habitat conservation and restoration efforts.

## Introduction

Land use change is causing dramatic resource declines for native wildlife (Gossner et al., 2016). Understanding how the availability of different resources impacts key ecosystem service providers is therefore crucial for informing conservation efforts (Davis et al., 2023; Requier & Leonhardt, 2020). This includes insect pollinators, which provide vital pollination services and are often threatened by habitat degradation (Potts et al., 2010). The importance of trophic resources for pollinators is firmly established, with higher flower abundance and diversity generally promoting bees (Hyjazie & Sargent, 2022; Kral-O’Brien et al., 2021). However, the roles of non-trophic resources such as shelter are largely not well understood (Davis et al., 2023; Requier & Leonhardt, 2020). Non-trophic resources can impact fitness as much as or more than trophic resources by affecting predation and pathogen rates, reproductive success, and interspecific interactions (e.g., Karban et al., 2013). Given a growing body of research that suggests non-trophic resources play critical roles in ecosystem function and resilience (Kéfi et al., 2015), there is an urgent need to understand how such resources alter pollinator communities. Understanding how land management impacts those resources is also imperative given widespread habitat loss and degradation (Scanes, 2018). Rangelands are a key context for this. These natural and semi-natural grasslands are managed through domestic livestock grazing that can dramatically alter trophic and non-trophic resources. Under appropriate management, they can also serve as natural habitat for wildlife (Havstad et al., 2007).

Shelter, including nests and overwintering sites, is a key non-trophic resource historically overlooked in pollinator research (Harmon-Threatt, 2020; Kremen et al., 2007). Common pollinator shelter locations include underground in soil and aboveground in living or dead vegetation, leaf litter, or cavities (Michener, 2007). Adults shelter for protection from heat, desiccation, and predation. For larvae that are largely immobile and spend significant time in nests (e.g. bees, many beetles), survival to adulthood relies heavily on appropriate nest site selection by the mother. Limited nesting habitat or poor nest site choice can leave offspring susceptible to predation, pathogens, floods, fires, or other natural disturbances (Antoine & Forrest, 2021). Availability of habitat with suitable shelter may therefore be a key driver of insect pollinator abundance, diversity, and community composition. This is particularly important for bees, which spend all but a short period of their life cycle inside a nest.

Over 80% of bee species nest in the ground, yet most investigations of ground-nesting bee habitat focus on aboveground vegetation cover and its correlate: bare ground (reviewed in Harmon-Threatt, 2020). Female bees searching for suitable nesting habitat may rapidly assess such characteristics. These characteristics can influence nest access and microclimate. Dense vegetation or leaf litter can protect nests from extreme weather (Song et al., 2013), and likely from nest parasites or predators (Antoine & Forrest, 2021). On the other hand, sparser cover can increase nest accessibility and facilitate nest location via visual or odor cues when bees return from foraging (Gomez Ramirez et al., 2023; Inouye, 2000; Ostwald et al., 2019; Wuellner, 1999). Bare ground also warms up more quickly in the spring and morning, potentially allowing bees to become active earlier in the year or day (Cane, 2021; Herb et al., 2008; Wuellner, 1999). While these mechanisms have not been explicitly tested, multiple studies have found higher bee abundance or diversity at sites with more bare ground (reviewed in Harmon-Threatt, 2020).

Belowground soil characteristics may be equally or more important for bee nesting and survival than aboveground shelter characteristics, but are rarely considered (Cane, 1991; Lybrand et al., 2020; Wuellner, 1999). This is in part due to challenges with locating underground nests. Habitat characteristics such as soil texture (proportions of sand, silt, and clay particles), compaction, and organic matter content can strongly affect nest conditions. Sandier or less compacted soils tend to be easier to dig in (Brockmann, 1979; Polidori et al., 2010; Potts & Willmer, 1997), and nests may have more structural integrity in more compacted soils (Potts & Willmer, 1997) or soils with more organic matter (Murphy, 2015). Rapid soil drainage (affected by texture, compaction, and organic matter content) may reduce flooding and fungal pathogens (Collis-George, 1991; Fellendorf et al., 2004; Murphy, 2015; Obayomi et al., 2019; Parr & Bertrand, 1960) or lead to brood cell desiccation (Potts and Willmer 1997). Documented ground-nesting bee preferences for sandier (Antoine, 2023) or less compacted soils (Sardiñas & Kremen, 2014) support these hypotheses. However, these habitat characteristics can also alter plant communities and cover (Dodd et al., 2002; Pennington et al., 2017; Renne et al., 2019). Thus, it is unclear whether documented bee responses to aboveground habitat characteristics indicate preferences for that characteristic itself, or for associated belowground habitat characteristics. There is a crucial need to determine impacts of belowground soil properties, in addition to aboveground vegetation, on bee communities.

These questions are particularly important to study in rangelands, where the distinction between suitable and unsuitable habitat is often less clear than in cropland or urban areas. Because they are semi-natural areas, rangelands not only generate farm revenue (Maher et al., 2021) but provide critical trophic and non-trophic resources for pollinators and other organisms in the 40-50% of Earth’s ice-free surface that they cover (Sala et al., 2017). However, unsustainable rangeland management practices (e.g., overgrazing) are degrading rangelands (Copeland et al., 2023; Goodwin et al., 2023). While low intensity or intermittent grazing can provide the disturbance necessary to promote flowering plants, overgrazing often reduces plant diversity (Apfelbaum et al., 2022; Barry & Huntsinger, 2021; DeBano et al., 2016). Grazing management can also alter habitat characteristics associated with ground-nesting bee shelter, for example by altering bare ground cover (Buckles & Harmon-Threatt, 2019; Shapira et al., 2020) or compacting soil (Kimoto et al., 2012). Despite rangelands’ high potential to support healthy pollinator communities, we still know relatively little about how grazing impacts pollinators (Hanberry et al., 2021). Studies to date have found positive, neutral, or negative effects of sustainability-oriented grazing practices (overviewed in Briske, 2017; T. Wang et al., 2020) on bee communities (reviewed in Kral-O’Brien et al., 2023; Thapa-Magar et al., 2022; Tonietto & Larkin, 2018). Understanding the mechanisms that drive these changes, including the importance of nesting resources in rangeland, is urgently needed (Kral-O’Brien et al., 2023; but see Buckles & Harmon-Threatt, 2019; Griffin et al., 2021; Shapira et al., 2020).

The present study focuses on ranches managed with adaptive multi-paddock grazing. This type of rotational grazing incorporates periods of rest by stocking grazers at high density in relatively small paddocks and frequently rotating them to a new paddock. It aims to enable plant regeneration and distribute livestock impacts across the entire landscape, rather than in a small number of preferred locations (Teague et al., 2011). Rotational grazing research, including a very limited body of adaptive multi-paddock grazing research, has largely focused on forage (mainly grasses) and soil impacts (reviewed in Byrnes et al., 2018). Of particular relevance for bees, adaptive multi-paddock grazing can increase vegetation biomass (Grissom & Steffens, 2013; Harmel et al., 2021), reduce bare ground (Teague et al., 2004), and increase floral richness (McDonald et al., 2019). Rotational grazing also may compact soil less than continuous grazing, although this has not been studied for adaptive multi-paddock grazing specifically (Byrnes et al., 2018). However, data are lacking on how adaptive multi-paddock grazing affects bees.

We addressed three key questions about how floral food and shelter impact bees in rangeland, focusing on several habitat characteristics that describe these resources. First, we asked how variation in food and shelter resources drive differences in ground-nesting bee assemblages. We hypothesized that nesting habitat characteristics are as important as floral abundance and diversity for bee assemblages because both trophic and non-trophic resources play key roles in bee survival. Second, we determined whether above- or belowground nesting habitat characteristics are more important determinants of ground-nesting bee assemblages. We hypothesized that belowground resources are stronger drivers of ground-nesting bee assemblages because these bees spend the majority of their lives, including the vulnerable egg and larval stages, underground. Third, we identified which habitat characteristics are sensitive to grazing. We hypothesized that areas with longer grazing-free “rest” periods provide better food for ground-nesting bees by allowing more forb growth. For ground-nesting bee shelter, longer rest periods may degrade aboveground nesting habitat by reducing bare ground, and conversely improve belowground nesting habitat by reducing soil compaction.

## Methods

### Study Sites

We conducted this study on ranches in the Cross Timbers ecoregion in North Texas (Fig. S1). The Cross Timbers are characterized by tallgrass prairies interspersed with dense woodland belts, with gently rolling hills. Precipitation is highly variable, averaging 686-965 mm per year, with two wet seasons: spring and fall. Summer temperatures are hot, averaging 21-36°C, and winter temperatures are mild at -1-3°C (Griffith et al., 2007). Soils range from heavy clays to clay-loams to loams (UC Davis Soil Resource Lab et al., 2022). Ranches were managed using adaptive multi-paddock grazing with cattle and sheep kept at high densities and rotated among small paddocks every 1-3 days. We sampled nine open grassland sites that were sufficiently far apart relative to insect flight distances to be spatially independent (separated by at least 1.5 km; Kendall et al., 2022). Sites additionally shared similar soil types based on soil maps (UC Davis Soil Resource Lab et al., 2022), although with sufficient variation to be able to produce differences in plant and bee communities. These sites were located on two management units, each of which contained a mixed herd of cattle and sheep that was rotated within the unit. Each site was in a separate paddock. We obtained grazing records to calculate grazing rest period (time since the site was last grazed by livestock) for each site prior to each sampling event. This rest period calculation was unique to each sampling season for each site.

At each site, we established eight 90 m transects radiating from the site’s center, one in each cardinal and intercardinal direction (Fig. S1). Along these transects we measured floral food, aboveground nesting habitat characteristics, and bee assemblages once during the spring (April-May), summer (June), and fall (September-October) bloom periods (three times per year) in 2021 and 2022 (six times total). We measured flowers and aboveground nesting habitat characteristics in six 1 × 1 m quadrats, located at 15 m intervals, along each transect (48 quadrats per site; Fig. S1). Our study region’s short bloom periods, and rainy and windy weather in spring and fall, limited suitable sampling days and restricted us to sampling once per bloom period. We measured belowground nesting habitat characteristics, which do not change substantially over short time periods (Zarekia et al., 2012), in spring 2021 (and soil compaction in spring 2022). We measured these characteristics at 30 and 90 m from the site center, walking 10 m to the left (30 m) or right (90 m) of the eight transects to avoid altering vegetation along transects (16 samples/site; Fig. S1). Table S1 describes these measurements. This study did not require ethical approval or permits.

### Floral Food

We measured each site’s floral abundance and richness (Table S1) by identifying each flowering plant to species (or occasionally genus) and counting the number of flowering units (individual flowers, umbels, or spikes depending on species, as in Lichtenberg et al., In press) of each plant species in each 1 × 1 m quadrat. To better capture floral species richness in heterogeneous grasslands, we recorded the presence of additional flowering species found within 1 m of each cardinal and intercardinal transect. At each site, we photographed each flowering plant species and confirmed identification with experts. Plant observations are available at https://www.inaturalist.org/projects/lichtenberg-lab-rotational-grazing-study.

### Aboveground Nesting Habitat Characteristics

In the same quadrats, we estimated spring, summer, and fall vegetation cover at both canopy (top of vegetation) and ground levels. Canopy cover types included live grass, live forbs, dead/senesced standing grass and forbs, wood (e.g., tree trunks), and no canopy. Ground cover types included litter, bare ground, rocks and gravel, moss, manure, green prostrate angiosperms, and stems. We quantified the portion of each quadrat’s 25 squares (each 20 × 20 cm), to the nearest half square, in each category then averaged cover proportions for each category for the site and season (as in Lichtenberg et al., In press).

### Belowground Nesting Habitat Characteristics

We measured several belowground habitat characteristics with potential to impact ground-nesting bee nest site selection or success: soil texture, depth, organic matter content, pH, and compaction (Table S1). We collected 30 cm long soil cores where the soil was deep enough, and several shorter cores with a total length of at least 30 cm in shallower soils. We mixed at least three cores at each sampling location to account for small-scale soil heterogeneity (Cunningham et al., 2012). The first complete core was used to determine maximum soil depth up to 30 cm. In the lab we measured soil texture, organic matter content, and pH of each sample using standard pipette analysis, loss on ignition (LOI), and soil pH procedures, respectively (Klute, 1986; Rousk et al., 2009). We measured soil compaction by collecting soil samples of consistent volume (90.59 cm^3^, using an AMS Bulk Density 2“ x 2“ SST Liner) and determining bulk density (a precise soil compaction proxy; Nawaz et al., 2013) using standard procedures (Klute, 1986).

### Bee Assemblages

We sampled each site’s bee assemblage via passive trapping and active netting on the same day as we measured aboveground vegetation in spring, summer, and fall (Table S1). Traps included one blue vane trap at the site’s center and four sets of three pan traps placed 5 m from the site’s center along each cardinal direction (Fig S1; 2 oz plastic bowls painted in colors attractive for bees: Rust-oleum Fluorescent yellow and white, Krylon bold neon blue; Kearns & Inouye, 1993; Leong & Thorp, 1999). We filled each trap with soapy water and deployed the traps for 24 hours. To reduce traps’ attraction of bees from long distances (Portman et al., 2020), we placed traps at vegetation level. At each site, we netted bees on flowers along transects at 10:00 and 13:00 for 30 minutes in 2021 and 45 minutes in 2022. Bees were only netted if they touched flower reproductive parts. We sampled when weather allowed for normal bee activity: fully or mostly sunny, temperatures above 18 °C, and average wind speed less than 6.7 m/s with gusts less than 8.9 m/s (as in Buckles & Harmon-Threatt, 2019; Stein et al., 2020). We later identified all specimens to genus or species using morphological characteristics and available keys (e.g., Bzdyk, 2012; Dumesh & Sheffield, 2012; Gardner & Gibbs, 2023; Michener et al., 1994; Portman et al., 2022; Williams et al., 2014; complete list in Table S2) and verified identifications with taxonomic experts. All specimens are stored in the Lichtenberg Lab and will ultimately be deposited in the University of North Texas Elm Fork Natural Heritage Museum.

### Data Analyses

We assessed the impacts of food and shelter resources on bee abundance, richness, and community composition. All analyses were conducted in R (R Core Team, 2023). First, we reviewed existing literature on bee nesting behavior to classify species as ground nesting if they nest in the ground obligately or facultatively, and nesting aboveground otherwise. Analyses classified both obligate and facultative ground-nesting bees as “ground nesters”. Next, we calculated ground-nesting bee abundance, richness, and Chao diversity (Chao et al., 2014) for each site and sampling season (e.g., summer 2021). Richness and Chao calculations excluded several specimens that could not be identified to species (e.g., damaged during collection) and thus could not be categorized as the same or different species from congeners at the same site. We also ensured that including specimens that were identified to the species group along with specimens that could be identified to the constituent species would not impact diversity calculations (i.e., they were not found at the same site in the same sampling season). Chao estimates were often very close to measured richness (Table S3; Spearman’s rank correlation: r = 0.94, S = 1394.97, p < 0.0001), indicating that our sampling effort was sufficient to accurately capture the bee assemblages at our sites, and thus we present analyses based on measured richness.

We determined the impacts of food and shelter resources on ground-nesting bee abundance and richness using (generalized) linear mixed models. These models included either bee abundance or richness as the response, habitat characteristics as fixed effects, and site and sampling season (which incorporated both year and time of year) as random effects. We used the identity link for the richness model (lme4 package; Bates et al., 2014) but a negative binomial model for bee abundance due to overdispersion (glmmTMB package; Brooks et al., 2017). Because quantitative predictors varied in units of measurement, we standardized predictors by subtracting the mean and dividing by the standard deviation.

We selected six fixed-effect predictors that were relevant to our hypotheses and captured a range of resource-associated characteristics: floral abundance, floral richness, bare ground cover, soil sand content, soil depth, and soil compaction (bulk density). We first eliminated habitat characteristics with insufficient variation for statistical analyses: wood, rocks and gravel, moss, manure, and green prostrate angiosperms cover; and soil organic matter content and pH. We also eliminated litter cover, which measures only litter surface area and not depth and thus is less informative in our system. Among the remaining variables, we considered which capture similar habitat characteristics. For example, flower abundance and richness closely relate to live forb cover but more directly measure food resources. Among these sets, we selected the variables that most directly measure the resources of interest or are most biologically relevant, are not inherently season dependent (e.g., more vegetation is senesced in fall), and are most commonly measured in similar studies (e.g., bare ground cover rather than no canopy cover). Initial analyses confirmed that our six predictors were not collinear (VIF < 4; Kroll & Song, 2013). Tables S4 and S5 show correlations among these variables and summarize them, respectively.

We determined how food and shelter affected ground-nesting bee community composition using analysis of multivariate data (Wang et al., 2012). First, we generated a matrix of the presence or absence of each species at each site in each year. We analyzed presence/absence data rather than abundances to prevent potential biases from social species. Analyses excluded specimens that could not be identified past genus and singleton species. In several cases (e.g., *Melissodes communis-tepaneca*) where some specimens within a species complex were identifiable to species and others to complex, we grouped all specimens as a single taxonomic entity. We summarized most habitat characteristics as the yearly median value across sampling events, and floral abundance and richness as the total number of flowering units or taxa, respectively, across all three seasons. We then used the mvabund R package (Wang et al., 2012) to fit a generalized linear model of multivariate data, with binomial family, that tested impacts of habitat characteristics on both overall community composition and each ground-nesting bee species. Analyses of impacts on individual species used Montecarlo resampling and adjusted for multiple sampling.

We determined grazing rest period impacts on food and shelter through a set of models with a habitat characteristic as the response, grazing rest period (summarized in Table S5) as a fixed effect, and season as a fixed effect and sampling season and site as random effects for characteristics measured seasonally (bare ground cover, floral abundance, floral richness). For habitat characteristics that can change with grazing on a very short timescale – floral abundance and richness, bare ground cover – we measured the grazing rest period from the last grazing event to the sampling date. For soil compaction, which likely responds to more than one cycle of grazing and rest, we measured the average grazing rest period in 2021: the year prior to bulk density measurement. We did not test impacts on soil sand content or soil depth, as geological processes such as severe erosion are the main forces that can change these properties (Dong et al., 2022). We used logistic regression for proportional response variables (bare ground cover) with proportions weighted by the number of squares within quadrats sampled per sampling event (1200), negative binomial regression for floral abundance, and the identity link for other variables. We conducted likelihood ratio tests to measure the relative effects of grazing rest period on each habitat characteristic.

## Results

We collected 1040 individual bees across 89 taxa and 30 genera. Ground nesters comprised 789 individuals (76% of all bees collected) spanning 74 taxa (83% of all taxa collected), 21 genera, and 6 families (Table S6). The majority of ground-nesting bees were in the family Halictidae (73%), followed by Apidae (24%). *Lasioglossum* species were the most common (60% of ground-nesting bees), followed by *Bombus* and *Melissodes* (each 11%), *Agapostemon* (6%), and *Halictus* (5%)(Fig. S2). Capture method did not affect which bee species we caught (Table S6).

Floral abundance affected bee abundance, but not bee richness (Table 1). Sites with more flowers had more ground-nesting bees (Fig. 1). However, floral richness did not affect ground-nesting bee abundance or richness (Table 1). Floral abundance and richness did not affect ground-nesting bee community composition (Table S7).

**Table 1:** Coefficients and likelihood ratio test statistics for the models investigating effects of food and shelter resources on ground-nesting bee abundance and richness. All predictors were standardized to facilitate comparison across habitat characteristics. Terms significant at α < 0.05 are highlighted in green.

| Bee community metric | Habitat characteristic | Coefficient | Coefficient Standard Error | $\chi^2$ | df | p-value |
| --- | --- | --- | --- | --- | --- | --- |
| Abundance | (Intercept) | 2.28 | 0.17 |  |  |  |
| Abundance | Floral abundance | 0.27 | 0.12 | 4.68 | 1 | 0.03 |
| Abundance | Floral richness | 0.13 | 0.17 | 0.56 | 1 | 0.45 |
| Abundance | Bare ground cover | -0.13 | 0.12 | 1.17 | 1 | 0.28 |
| Abundance | Soil sand content | 0.89 | 0.19 | 16.77 | 1 | <0.0001 |
| Abundance | Soil depth | -0.06 | 0.15 | 0.16 | 1 | 0.69 |
| Abundance | Soil compaction (bulk density) | -0.55 | 0.16 | 10.13 | 1 | 0.002 |
| Richness | (Intercept) | 5.32 | 0.81 |  |  |  |
| Richness | Floral abundance | 0.67 | 0.52 | 1.60 | 1 | 0.21 |
| Richness | Floral richness | 0.32 | 0.77 | 0.17 | 1 | 0.68 |
| Richness | Bare ground cover | 0.13 | 0.53 | 0.07 | 1 | 0.80 |
| Richness | Soil sand content | 2.02 | 0.76 | 6.39 | 1 | 0.01 |
| Richness | Soil depth | 0.12 | 0.59 | 0.04 | 1 | 0.84 |
| Richness | Soil compaction (bulk density) | -1.39 | 0.68 | 3.89 | 1 | 0.049 |

**Figure 1:**
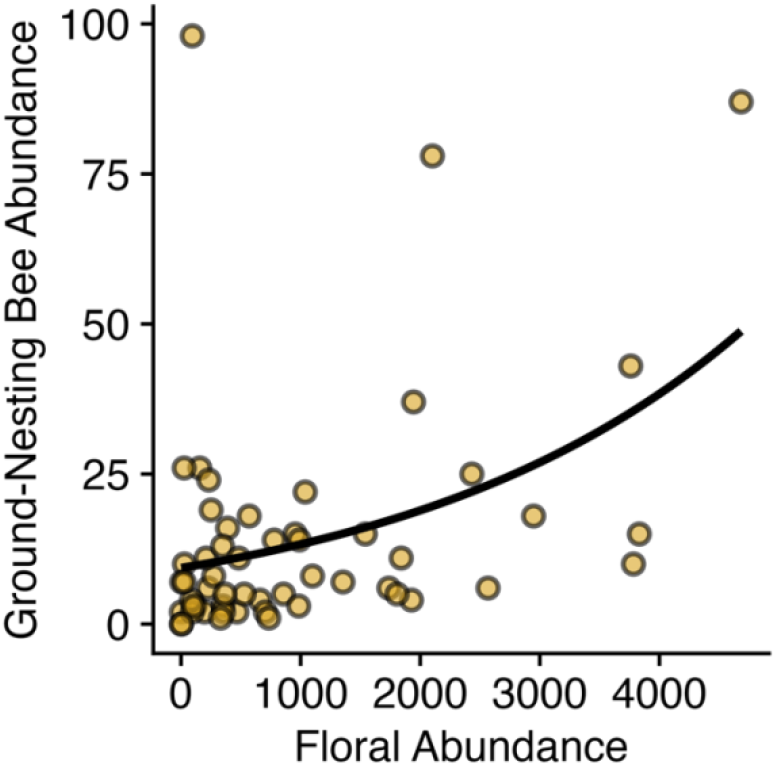
Sites with more flowers had more ground-nesting bees within a sampling season. Floral abundance indicates the total number of flowering units counted in 48 1 × 1 m quadrats at a site within a season. The curve indicates best fit.

Some belowground nesting habitat characteristics, but no aboveground nesting habitat characteristics, affected bee assemblages (Table 1). Sandier and less compacted (lower bulk density) soils supported higher ground-nesting bee abundance and richness (Fig. 2). However, soil depth and bare ground cover did not impact ground-nesting bee abundance or richness. No nesting habitat characteristics—including soil sand content, soil compaction, soil depth, and bare ground cover—altered ground-nesting bee community composition (Table S7).

**Figure 2:**
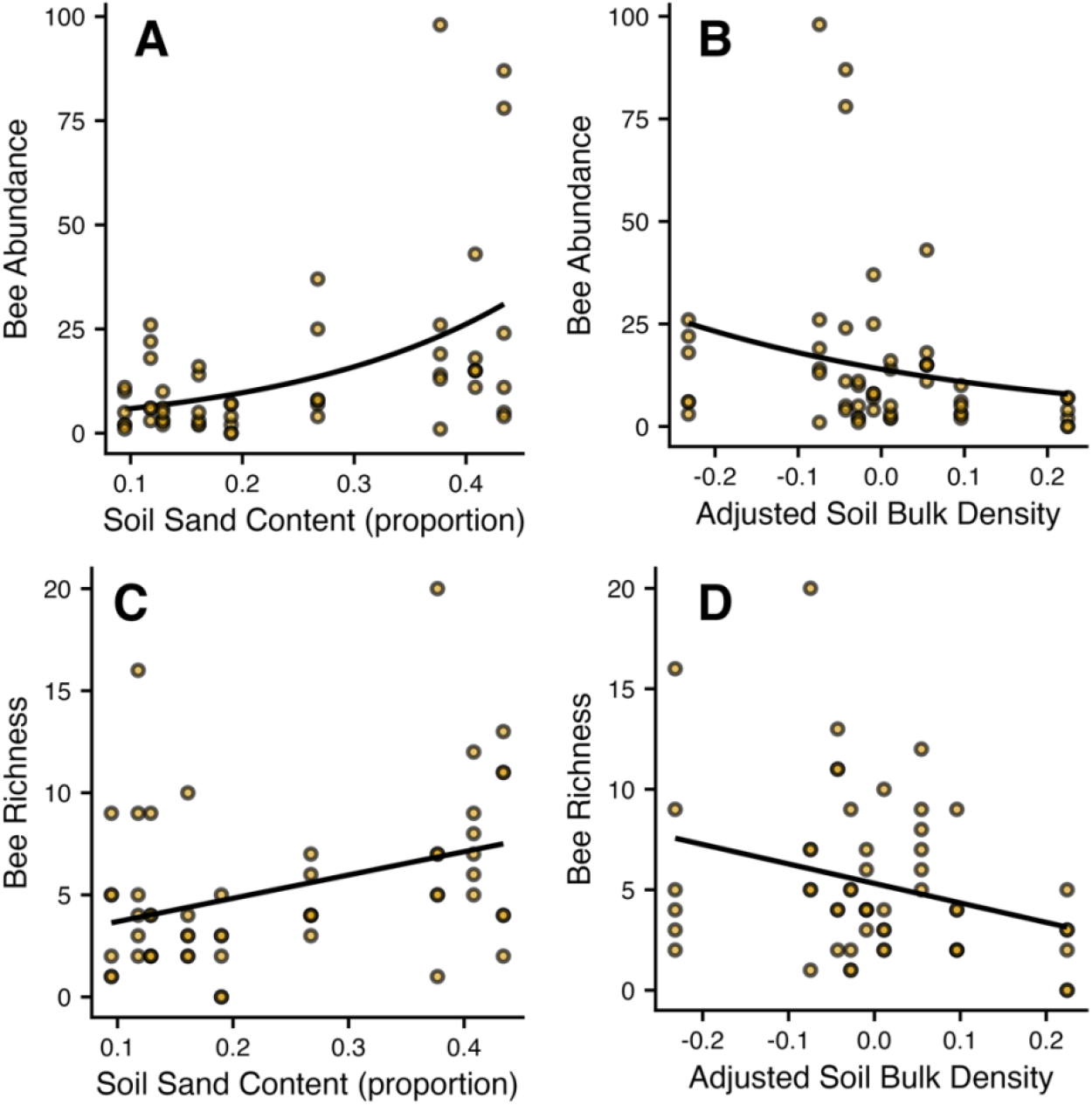
Ground-nesting bee (A, B) abundance and (C, D) richness were (A, C) higher at sites with sandier soils and (B, D) lower at sites with more compacted soils. B and D are marginal effects plots of bee abundance or richness against the residuals of soil bulk density, controlling for soil sand content. Curves and lines indicate best fit.

Sites with longer grazing rest periods had higher floral abundance and richness, and more bare ground (Fig. 3; Table S8). Thus, grazing may indirectly impact ground-nesting bee abundance through impacts on floral abundance. Additionally, floral richness and bare ground cover were lower in fall than in spring or summer (Table S8).

**Figure 3:**
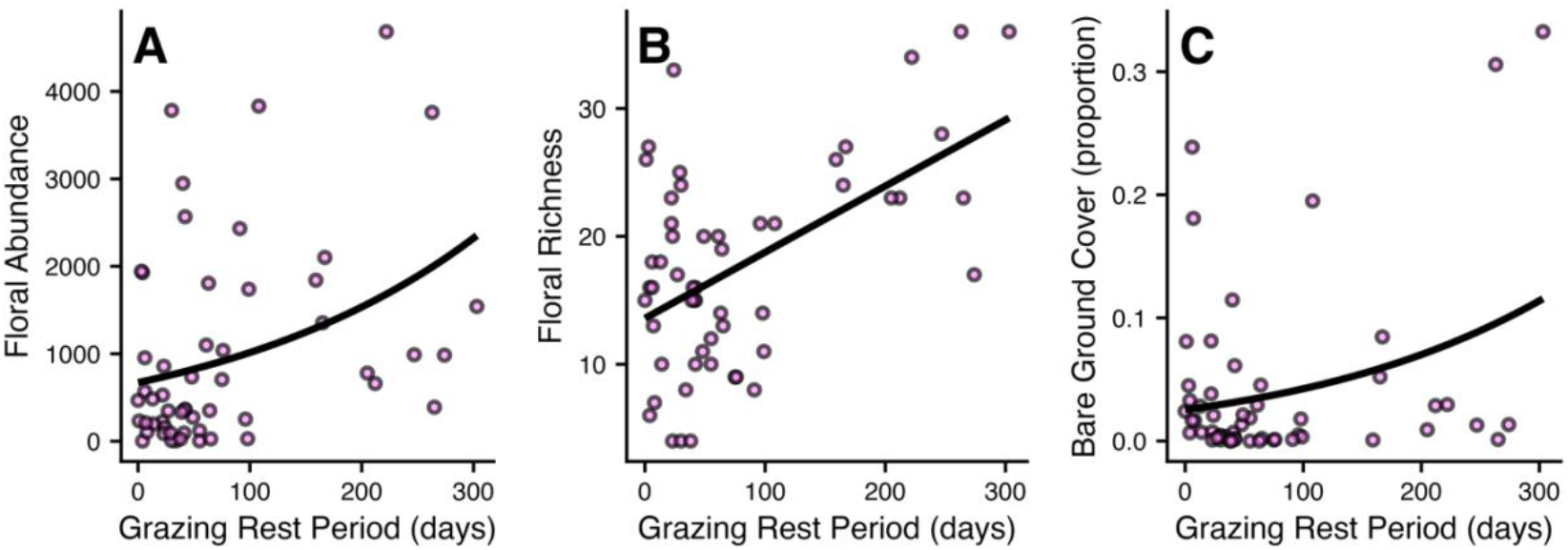
Sites with longer grazing rest periods had more (A) flowers, (B) flower species, and (C) bare ground. Curves and line indicate best fit.

## Discussion

Determining the impacts of trophic and non-trophic resources is critical for effective rangeland management for pollinator conservation. Floral food availability often promotes pollinators (Potts et al., 2003; Prendergast et al., 2022), and is sometimes assumed to be the main resource limiting bee communities. Contrary to this, but consistent with several recent studies (Antoine, 2023; Brokaw et al., 2023; Wersebeckmann et al., 2023), we found that belowground nesting habitat characteristics predicted ground-nesting bee assemblages as well as or better than floral food resources. Ground-nesting bee abundance was highest at sites with sandier and less compacted soils, and with more flowers. Ground-nesting bee richness was higher at sites with sandier and less compacted soil. Overall, our results demonstrate that belowground habitat characteristics may strongly influence bee nesting habitat availability.

We hypothesized that soil texture is important for ground-nesting bee assemblages because sandier soils may be easier to dig in (Potts and Willmer, 1997) and reduce nest flooding due to greater water permeability (Fellendorf et al., 2004). We found that ground-nesting bees were indeed more abundant and speciose in sandier soil. These findings are consistent with observational studies of bee nesting habitat (Antoine, 2023; Cane, 1991). However, previous experimental manipulations of soil texture found no difference in nest abundance across soils (Antoine, 2023; Fortel et al., 2016; Tsiolis et al., 2022). The range of soil textures available to bees in each study may explain this discrepancy, as the soils in most experiments were at least 50% sand. In contrast, our sites had relatively low sand content (10-43%, Table S5), ranging from clay-loams to heavy clays. Approximately 35% of topsoil worldwide contains similarly low sand percentages (Shangguan et al., 2014), including humid areas (He et al., 2024) and many grasslands (Shangguan et al., 2014). Thus, our results suggest that predominance of an extreme soil type such as clay can restrict ground nesting niche space and limit bee nesting (Neff & Simpson, 1997; Strickler, 1979). Under such conditions, even small differences in soil sand content may noticeably benefit ground-nesting bees (as in Leone et al., 2022).

We also hypothesized that ground-nesting bees prefer less compacted soils, which are generally easier to dig in and have better drainage. Supporting this, we found that ground-nesting bees were more abundant and diverse in less compacted (lower bulk density) soils. Because we included both soil bulk density and sand content in our analyses, our results indicate that ground-nesting bees respond to soil compaction itself. Our results are consistent with a previous study indicating that higher soil compaction reduces ground-nesting bee abundance (Sardiñas & Kremen, 2014), but inconsistent with two that found no effect (Antoine, 2023; Tsiolis et al., 2022). However, comparing across studies is complicated by differences in soil compaction proxies. Most pollinator studies measured soil penetration resistance, which is more influenced by soil moisture and user technique and thus is perhaps less suitable for studies over time spans or in regions such as ours with frequent soil moisture fluctuation. We used bulk density, which is more precise (Vomocil, 1957). Additionally, soil compaction may be a particularly important factor determining ground-nesting bee habitat in regions with clayey soils, as clay soils have more total pore space than sandy soils (Singh et al., 2016) and thus are more susceptible to soil compaction despite having lower bulk density (Nawaz et al., 2013). Experimentally manipulating soil compaction while controlling potentially confounding factors such as soil texture, soil moisture, and floral abundance is necessary to determine when and the extent to which soil compaction directly reduces bee abundance and diversity. Furthermore, because soil substrate preferences can vary widely across species (Potts & Willmer, 1997; Wuellner, 1999), studies that incorporate bee functional traits (such as Aguirre-Gutiérrez et al., 2016) will allow for more nuanced assessments of ground-nesting bee nesting habitat.

In contrast to our results, bare ground has often been found to promote ground-nesting bees (reviewed in Harmon-Threatt, 2020). Such studies, which typically focus on community assembly or community-level responses to anthropogenic impacts, rarely measure soil texture or other belowground shelter characteristics and thus may attribute belowground drivers of community assembly to aboveground measures such as bare ground. Soil texture can strongly influence vegetative growth and thus bare ground cover (Augustine et al., 2020; Renne et al., 2019). Consistent with this, we found more bare ground at sites with sandier soils (Table S4; habitat characteristics were not collinear). As discussed above, studies investigating the natural history or population ecology of individual ground-nesting bee species have documented preferences for soils with specific belowground properties (e.g., several species studied by Antoine, 2023 and others cited in their Table 2-S1; Cane, 1991; López-Uribe et al., 2015; Potts & Willmer, 1997). Our results highlight the pressing need to integrate population and community ecology approaches by including belowground characteristics of shelter habitat when measuring resource availability for bees and other insects that shelter below ground.

A growing body of literature aims to understand how grazing lands can support pollinator conservation (synthesized in Kral-O’Brien et al., 2023; Thapa-Magar et al., 2022; Tonietto & Larkin, 2018). This goal is complicated by large variation in grazing practices and by responses to management that depend on factors such as aridity (Thapa-Magar et al., 2022), habitat type (Elwell et al., 2016; Kohler et al., 2020), and landscape composition (Griffin et al., 2021). Better understanding the mechanisms through which rangeland management impacts ground-nesting bee communities is critical for sustainable rangeland management. We thus also investigated which bee food- and nest-associated habitat characteristics are sensitive to grazing rest period. Sites with longer grazing rest periods had higher floral abundance and richness, and more bare ground. This is consistent with research showing that rotational grazing can promote plant diversity or flower availability (Harmel et al., 2021; McDonald et al., 2019; Ravetto Enri et al., 2017). Combined with our bee abundance analyses, this suggests that adaptive multi-paddock grazing may help increase ground-nesting bee abundance by promoting flowering forbs. A similar mechanism has been proposed with moderate grazing intensity (Lasway et al., 2022; Lázaro et al., 2016; Vulliamy et al., 2006), other rotational grazing schemes (Mitchell et al., 2023; Ravetto Enri et al., 2017), and grazing for grassland restoration (Stein et al., 2020) or maintenance (Veen et al., 2024).

Our study highlights the importance of considering non-trophic resources when managing for biodiversity conservation. This includes soil properties for species that live belowground. We found that a commonly-used aboveground proxy of bee nesting habitat, bare ground, may be inadequate for assessing bee habitat quality. Land managers should use this measure with caution until further research can provide guidelines on the contexts in which bare ground is a useful habitat indicator. Moreover, belowground nesting habitat characteristics may be particularly important for ground-nesting bee conservation in areas with relatively high floral availability (Griffin et al., 2021), such as rangelands. This could necessitate managing for ground-nesting bees’ preferred levels of soil compaction in a given soil type, for example by using an optimal grazing intensity (Daniel et al., 2002; Kimoto et al., 2012). Habitat characteristics like soil texture that are generally unaffected by land management may also strongly regulate bee assemblages. Thus, conservation and restoration planning should consider not only which management actions or types to implement, but also how management-insensitive habitat characteristics like soil texture can inform site prioritization. Overall, more ecological research on working lands that (1) measures belowground nesting habitat characteristics and (2) is conducted across diverse soil types is urgently needed to improve evidence-based recommendations for pollinator conservation and restoration efforts.

## Acknowledgements

We thank the Dixon Water Foundation for ranch access; Alejandra Gage, Christopher Graffam, Alyssa Neathery, John Linogao, Pablo Lopez, Brandon Meadows, Marie Muñiz, Grace Phi, Brand Richter, and Michelle Vohs for field and lab assistance; Jack Neff and Nick Medina for bee identification; Isaac Eastland and Brand Richter for plant identification; Zacchaeus Compson for pH meter access; and Alex Harmon-Threatt, David Hoeinghaus, and three anonymous reviewers for feedback. This project was funded by a Garden Club of America Board of Associates Centennial Pollinator Fellowship to SMC and a College of Science Seed Grant to EML.

## Data availability

Data and R scripts available from the Zenodo repository https://zenodo.org/records/15258813.

