## Supplementary material for "Local nesting resources rather than floral food alter ground-nesting bee communities in grazed habitats": Table S

**Table S1:** Types of habitat and bee assemblage data collected, including with how they were measured and with what frequency.

| **Measured in field or lab?** | **Property** | **Sampling** | **Measurement** | **Frequency** |
|---|---|---|---|---|
| Field | Flowers | Sampled at six locations every 15 m along each transect | Floral abundance and richness per site (all quadrats combined) | 3x/year |
| Field | Vegetation cover | Sampled at six locations every 15 m along each transect | Using quadrats (1 $m^2$) split into 25 even squares, cover types counted by halves of squares (2% resolution) at both canopy and ground levels | 3x/year |
| Lab | Soil texture | 4+ soil cores taken at each sampling location and mixed | Standard pipette analysis | Once (2021) |
| Field | Soil depth | One complete soil core, sampled at 30 m and 90 m on each transect | Maximum core length, up to 30 cm | Once (2021) |
| Lab | Soil organic matter | Same as sampling for soil texture | Standard LOI analysis and conversion of organic carbon to organic matter content | Once (2021) |
| Lab | Soil pH | Same as sampling for soil texture | Standard soil pH analysis with pH probe, 2:1 soaked and suspended solution | Once (2021) |
| Lab | Soil compaction | Soil samples collected by evenly hammering a 5.08 $cm^2$ stainless steel liner | Standard bulk density analysis | Once (2022) |
| Field | Bee assemblages | One blue vane trap; four sets (3 colors each) of pan traps; 60 (2021) or 90 (2022) minutes of netting bees on flowers | Bee abundance, richness, community composition | 3x/year |
| Lab | Grazing rest period | Determined for each site and each sampling event | Number of days since the site was last grazed by livestock | 3x/year |

**Table S2:** Taxonomic resources used to identify insects.

| **Taxonomic group** | **Source(s)** |
|---|---|
| Anthophila (bees) | (Michener et al., 1994; Mitchell, 1960, 1962) |
| *Agapostemon* | (Roberts, 1972) |
| *Andrena* | (Bouseman & LaBerge, 1978; B. N. Danforth, 1994; Donovan, 1977; La Berge, 1973; W. LaBerge, 1967; W. E. LaBerge, 1969, 1971, 1977, 1980, 1985, 1986, 1989; W. E. LaBerge & Bouseman, 1970; W. E. LaBerge & Ribble, 1972, 1975; W. E. LaBerge & Thorp, 2006) |
| *Anthidiellum* | (T. L. Griswold & Michener, 1988) |
| *Anthophora* | (Brooks, 1983) |
| *Atoposmia* | (Hurd & Michener, 1955; Michener, 1939) |
| *Augochlorella* | (Coelho, 2004) |
| *Bombus* | (P. H. Williams et al., 2014) |
| *Calliopsis* | (Shinn, 1967) |
| *Centris* | (Snelling, 1974; Vivallo, 2013, 2020) |
| *Ceratina* | (Daly, 1973) |
| *Colletes* | (Stephens, 1954; P. H. Timberlake, 1951) |
| *Dieunomia* | (Blair & Cockerell, 1935) |
| *Dufourea* | (Dumesh & Sheffield, 2012) |
| *Epeolus* | (Onuferko, 2018) |
| *Epimelissodes* | (W. E. LaBerge, 1958) |
| *Eucera* | (P. H. Timberlake, 1969) |
| *Halictus* | (Roberts, 1973; P. Timberlake, 1961) |
| *Heriades* | (Michener, 1938) |
| *Hesperapis* | (Stage, 1966) |
| *Hylaeus* | (Snelling, 1970) |
| *Lasioglossum* | (Gardner & Gibbs, 2020, 2023; Gibbs et al., 2013) |
| *Lithurgopsis* | (Mitchell, 1938) |
| *Megachile* (*Litomegachile*) | (Bzdyk, 2012; Gonzalez & Griswold, 2007) |
| *Melissodes* | (W. LaBerge, 1957, 1961; W. E. LaBerge, 1956) |
| *Melitoma* | (Krombein et al., 1979; Schuh et al., 2010) |
| *Osmia* | (T. Griswold & Rightmyer, 2017; Rightmyer et al., 2010) |
| *Panurginus* | (Crawford, 1926) |
| *Perdita* | (B. Danforth, 1996; T. Griswold & Miller, 2010; Z. Portman et al., 2016; Z. M. Portman & Griswold, 2017; Snelling & Danforth, 1992) |
| *Protoxea* | (Hurd & Linsley, 1976) |
| *Stelis* | (Parker & Bohart, 1979) |
| *Trachusa* | (Brooks & Griswold, 1988) |
| *Xenoglossa* | (Ayala & Griswold, 2012) |
| *Xylocopa* | (Hurd, 1961) |

**Table S3:** Measured and estimated (Chao) ground-nesting bee richness at each site.

| Site | Sampling season | Measured ground-nesting bee richness | Estimated ground-nesting bee richness |
|---|---|---|---|
| BLO | spring 2021 | 8 | 10 |
| CRA | spring 2021 | 5 | 8 |
| FSO | spring 2021 | 2 | 2 |
| FTO | spring 2021 | 4 | 4.5 |
| HBD | spring 2021 | 2 | 3 |
| LAA | spring 2021 | 5 | 5.5 |
| LMP | spring 2021 | 13 | 16.3 |
| PMP | spring 2021 | 2 | 3 |
| PSO | spring 2021 | 2 | 3 |
| BLO | summer 2021 | 9 | 14 |
| CRA | summer 2021 | 2 | 2 |
| FSO | summer 2021 | 4 | 7 |
| FTO | summer 2021 | 4 | 10 |
| HBD | summer 2021 | 0 | |
| LAA | summer 2021 | 7 | 9 |
| LMP | summer 2021 | 4 | 5.5 |
| PMP | summer 2021 | 2 | 3 |
| PSO | summer 2021 | 5 | 15 |
| BLO | fall 2021 | 7 | 10 |
| CRA | fall 2021 | 4 | 7 |
| FSO | fall 2021 | 2 | 3 |
| FTO | fall 2021 | 3 | 4 |
| HBD | fall 2021 | 3 | 3.5 |
| LAA | fall 2021 | 1 | 1 |
| LMP | fall 2021 | 2 | 2 |
| PMP | fall 2021 | 3 | 6 |
| PSO | fall 2021 | 1 | 1 |
| BLO | spring 2022 | 12 | 17 |
| CRA | spring 2022 | 3 | 4 |
| FSO | spring 2022 | 2 | 2 |
| FTO | spring 2022 | 7 | 7.3 |
| HBD | spring 2022 | 0 | |

| **Site** | **Sampling season** | **Measured ground-nesting bee richness** | **Estimated ground-nesting bee richness** |
|---|---|---|---|
| LAA | spring 2022 | 7 | 12 |
| LMP | spring 2022 | 11 | 17 |
| PMP | spring 2022 | 4 | 5 |
| PSO | spring 2022 | 9 | 23 |
| BLO | summer 2022 | 6 | 7.5 |
| CRA | summer 2022 | 16 | 23.5 |
| FSO | summer 2022 | 9 | 45 |
| FTO | summer 2022 | 4 | 4 |
| HBD | summer 2022 | 5 | 6 |
| LAA | summer 2022 | 20 | 36.5 |
| LMP | summer 2022 | 11 | 13.5 |
| PMP | summer 2022 | 10 | 20.5 |
| PSO | summer 2022 | 5 | 5.5 |
| BLO | fall 2022 | 5 | 5.5 |
| CRA | fall 2022 | 9 | 14 |
| FSO | fall 2022 | 4 | 5.5 |
| FTO | fall 2022 | 6 | 8 |
| HBD | fall 2022 | 3 | 3 |
| LAA | fall 2022 | 5 | 6.5 |
| LMP | fall 2022 | 4 | 7 |
| PMP | fall 2022 | 3 | 4 |
| PSO | fall 2022 | 1 | 1 |

**Table S4:** Correlations among habitat characteristics, determined via Spearman's rank correlation tests. We include characteristics used in regressions and community composition analyses, as well as those that were excluded due to lower biological relevance or high dependence on season. We do not include characteristics with insufficient variation for statistical analyses (e.g., rocky ground cover).

| Habitat characteristic 1 | Habitat characteristic 2 | Test statistic (S) | r | p-value |
|---|---|---|---|---|
| Floral abundance | Soil compaction | 25,245.00 | 0.04 | 0.7865 |
| Floral abundance | Soil depth | 36,376.45 | -0.39 | 0.0039 |
| Floral abundance | Soil sand content | 20,089.76 | 0.23 | 0.0882 |
| Floral abundance | Bare ground cover | 16,203.92 | 0.38 | 0.0043 |
| Floral abundance | Live forb canopy cover | 10,026.26 | 0.62 | <0.0001 |
| Floral abundance | Live grass canopy cover | 33,444.41 | -0.27 | 0.0443 |
| Floral abundance | No canopy cover | 21,892.75 | 0.17 | 0.2317 |
| Floral abundance | Senesced vegetation canopy cover | 29,934.78 | -0.14 | 0.3091 |
| Floral abundance | Stem ground cover | 30,274.46 | -0.15 | 0.2663 |
| Floral abundance | Grazing rest period | 16,504.77 | 0.37 | 0.0058 |
| Floral richness | Soil compaction | 22,857.01 | 0.13 | 0.3535 |
| Floral richness | Soil depth | 34,380.85 | -0.31 | 0.0223 |
| Floral richness | Soil sand content | 15,514.87 | 0.41 | 0.0022 |
| Floral richness | Bare ground cover | 12,079.18 | 0.54 | <0.0001 |
| Floral richness | Live forb canopy cover | 10,285.06 | 0.61 | <0.0001 |
| Floral richness | Live grass canopy cover | 30,431.64 | -0.16 | 0.2479 |
| Floral richness | No canopy cover | 15,074.16 | 0.43 | 0.0013 |
| Floral richness | Senesced vegetation canopy cover | 41,012.12 | -0.56 | <0.0001 |
| Floral richness | Stem ground cover | 31,403.51 | -0.20 | 0.1533 |
| Floral richness | Grazing rest period | 18,728.91 | 0.29 | 0.0360 |
| Soil compaction | Soil depth | 42,413.25 | -0.62 | <0.0001 |
| Soil compaction | Soil sand content | 4,372.50 | 0.83 | <0.0001 |
| Soil compaction | Bare ground cover | 17,150.27 | 0.35 | 0.0103 |
| Soil compaction | Live forb canopy cover | 26,494.57 | -0.01 | 0.9434 |
| Soil compaction | Live grass canopy cover | 29,355.85 | -0.12 | 0.3916 |
| Soil compaction | No canopy cover | 20,965.17 | 0.20 | 0.1453 |
| Soil compaction | Senesced vegetation canopy cover | 31,329.87 | -0.19 | 0.1594 |
| Soil compaction | Stem ground cover | 26,325.55 | 0.00 | 0.9802 |
| Soil compaction | Grazing rest period | 28,523.21 | -0.09 | 0.5306 |
| Soil depth | Soil sand content | 41,538.75 | -0.58 | <0.0001 |

| Habitat characteristic 1 | Habitat characteristic 2 | Test statistic (S) | r | p-value |
|---|---|---|---|---|
| Soil depth | Bare ground cover | 39,632.56 | -0.51 | <0.0001 |
| Soil depth | Live forb canopy cover | 38,308.16 | -0.46 | 0.0005 |
| Soil depth | Live grass canopy cover | 16,057.52 | 0.39 | 0.0037 |
| Soil depth | No canopy cover | 33,195.04 | -0.27 | 0.0525 |
| Soil depth | Senesced vegetation canopy cover | 21,520.43 | 0.18 | 0.1935 |
| Soil depth | Stem ground cover | 24,230.78 | 0.08 | 0.5830 |
| Soil depth | Grazing rest period | 29,779.01 | -0.14 | 0.3301 |
| Soil sand content | Bare ground cover | 12,124.68 | 0.54 | <0.0001 |
| Soil sand content | Live forb canopy cover | 20,892.63 | 0.20 | 0.1397 |
| Soil sand content | Live grass canopy cover | 32,772.49 | -0.25 | 0.0692 |
| Soil sand content | No canopy cover | 18,526.44 | 0.29 | 0.0310 |
| Soil sand content | Senesced vegetation canopy cover | 31,299.69 | -0.19 | 0.1619 |
| Soil sand content | Stem ground cover | 30,394.36 | -0.16 | 0.2522 |
| Soil sand content | Grazing rest period | 27,641.74 | -0.05 | 0.7002 |
| Bare ground cover | Live forb canopy cover | 15,918.72 | 0.39 | 0.0033 |
| Bare ground cover | Live grass canopy cover | 35,917.15 | -0.37 | 0.0060 |
| Bare ground cover | No canopy cover | 18,512.70 | 0.29 | 0.0307 |
| Bare ground cover | Senesced vegetation canopy cover | 29,519.32 | -0.13 | 0.3671 |
| Bare ground cover | Stem ground cover | 22,514.23 | 0.14 | 0.3063 |
| Bare ground cover | Grazing rest period | 27,857.39 | -0.06 | 0.6569 |
| Live forb canopy cover | Live grass canopy cover | 30,131.22 | -0.15 | 0.2838 |
| Live forb canopy cover | No canopy cover | 24,169.88 | 0.08 | 0.5715 |
| Live forb canopy cover | Senesced vegetation canopy cover | 35,126.68 | -0.34 | 0.0122 |
| Live forb canopy cover | Stem ground cover | 23,752.72 | 0.09 | 0.4962 |
| Live forb canopy cover | Grazing rest period | 16,114.69 | 0.39 | 0.0040 |
| Live grass canopy cover | No canopy cover | 39,917.52 | -0.52 | <0.0001 |
| Live grass canopy cover | Senesced vegetation canopy cover | 32,207.34 | -0.23 | 0.0978 |
| Live grass canopy cover | Stem ground cover | 21,452.54 | 0.18 | 0.1871 |
| Live grass canopy cover | Grazing rest period | 28,347.94 | -0.08 | 0.5626 |
| No canopy cover | Senesced vegetation canopy cover | 40,025.29 | -0.53 | <0.0001 |
| No canopy cover | Stem ground cover | 37,590.08 | -0.43 | 0.0011 |
| No canopy cover | Grazing rest period | 26,095.97 | 0.01 | 0.9697 |

| Habitat characteristic 1 | Habitat characteristic 2 | Test statistic (S) | r | p-value |
|---|---|---|---|---|
| Senesced vegetation canopy cover | Stem ground cover | 20,382.33 | 0.22 | 0.1049 |
| Senesced vegetation canopy cover | Grazing rest period | 30,282.43 | -0.15 | 0.2653 |
| Stem ground cover | Grazing rest period | 30,039.52 | -0.15 | 0.2954 |

**Table S5:** Summary of habitat characteristics that describe food and shelter resources. Grazing rest period summarizes individual seasons, and not the site averages used in the analysis of grazing impacts on soil compaction.

| **Habitat characteristic** | **Minimum value** | **Median value** | **Maximum value** |
|---|---|---|---|
| Floral abundance (# flowering units) | 2 | 505.5 | 4683 |
| Floral richness (# species) | 4 | 16.5 | 36 |
| Bare ground cover (proportion) | 0.0 | 0.01 | 0.33 |
| Soil sand content (proportion) | 0.095 | 0.190 | 0.434 |
| Soil depth | 17.7 | 22.4 | 28.9 |
| Soil compaction (bulk density, $g/cm^3$) | 1.15 | 1.51 | 1.69 |
| Grazing rest period (days) | 0 | 55 | 303 |

**Table S6:** Nest location and total abundance of each taxon we collected. Morphospecies names are unique taxa that either cannot be reliably distinguished from each other (e.g., "*angelicus-texanus*" or "*metallica* complex"), or have not yet been formally described (e.g., "sp1A"). Names follow those used by regional taxonomic experts. Nest location is "Belowground" for species with underground nests (including those that nest above and below ground), and "Aboveground" for species that nest only above the soil surface. The source column provides nest location references. Specimens that could be identified only to family or genus were not classified or included in analyses.

| **Family** | **Nest Location** | **Source** | **Genus** | **Species** | **Abundance** | **Collection Method(s)** |
|---|---|---|---|---|---|---|
| Andrenidae | Belowground | (Rozen, 2018) | *Protoxaea* | *gloriosa* | 1 | Vane trap |
| Apidae | Belowground | (Michener, 2007) | *Anthophora* | *affabilis* | 1 | Vane trap |
| Apidae | Belowground | (Michener, 2007) | *Anthophora* | *dammersi* | 1 | Vane trap |
| Apidae | Belowground | (Orr, 2020) | *Anthophora* | *phenax* | 4 | Net, pan trap |
| Apidae | Aboveground | (Seeley & Morse, 1976) | *Apis* | *mellifera* | 172 | Net, pan trap, vane trap |
| Apidae | Belowground | (P. H. Williams et al., 2014) | *Bombus* | *fraternus* | 1 | Net |
| Apidae | Belowground | (P. H. Williams et al., 2014) | *Bombus* | *griseocollis* | 2 | Net |
| Apidae | Belowground | (P. H. Williams et al., 2014) | *Bombus* | *pensylvanicus* | 80 | Net, pan trap, vane trap |
| Apidae | Belowground | (P. H. Williams et al., 2014) | *Bombus* | sp. | 1 | Net |
| Apidae | Belowground | (Alcock et al., 1976) | *Centris* | *cockerelli* | 1 | Net |
| Apidae | Aboveground | (Rau, 1928; Rehan & Richards, 2010) | *Ceratina* | spp. | 32 | Net, pan trap, vane trap |
| Apidae | Belowground | (Ordway, 1987) | *Diadasia* | *rinconis* | 1 | Vane trap |

| **Family** | **Nest Location** | **Source** | **Genus** | **Species** | **Abundance** | **Collection Method(s)** |
|---|---|---|---|---|---|---|
| Apidae | Belowground | (Onuferko, 2017) | *Epeolus* | *lectoides* | 1 | Net |
| Apidae | Belowground | (Carril & Wilson, 2021) | *Epimelissodes* | *aegis* | 1 | Net |
| Apidae | Belowground | (Carril & Wilson, 2021) | *Epimelissodes* | *obliquus* | 1 | Net |
| Apidae | Belowground | (Scott et al., 2011) | *Eucera* | *belfragei* | 2 | Vane trap |
| Apidae | Belowground | (Scott et al., 2011) | *Eucera* | *speciosa* | 3 | Trap (type not recorded) |
| Apidae | Belowground | (Parker et al., 1981) | *Melissodes* | *agilis* | 7 | Net, vane trap |
| Apidae | Belowground | (B. N. Danforth et al., 2019) | *Melissodes* | *boltoniae-tinctus* | 2 | Net, trap (type not recorded) |
| Apidae | Belowground | (B. N. Danforth et al., 2019) | *Melissodes* | *communis* | 23 | Net, pan trap, vane trap |
| Apidae | Belowground | (B. N. Danforth et al., 2019) | *Melissodes* | *communis-tepaneca* | 3 | Vane trap |
| Apidae | Belowground | (B. N. Danforth et al., 2019) | *Melissodes* | *comptoides* | 3 | Pan trap, vane trap |
| Apidae | Belowground | (B. N. Danforth et al., 2019) | *Melissodes* | *coreopsis* | 24 | Net, vane trap |
| Apidae | Belowground | (B. N. Danforth et al., 2019) | *Melissodes* | *menuachus* | 1 | Vane trap |
| Apidae | Belowground | (B. N. Danforth et al., 2019) | *Melissodes* | *rivalis* | 1 | Vane trap |
| Apidae | Belowground | (B. N. Danforth et al., 2019) | *Melissodes* | *tepaneca* | 16 | Pan trap, vane trap |
| Apidae | Belowground | (B. N. Danforth et al., 2019) | *Melissodes* | *tinctus* | 1 | Pan trap |

| Family | Nest Location | Source | Genus | Species | Abundance | Collection Method(s) |
|---|---|---|---|---|---|---|
| Apidae | Belowground | (B. N. Danforth et al., 2019) | *Melissodes* | *vernoniae* | 2 | Net, vane trap |
| Apidae | Belowground | (B. N. Danforth et al., 2019) | *Melitoma* | *grisella* | 1 | Vane trap |
| Apidae | Belowground | (Hurd et al., 1974) | *Xenoglossa* | *pruinosa* | 2 | Net |
| Apidae | Aboveground | (Balduf, 1962) | *Xylocopa* | *virginica* | 21 | Net, pan trap, vane trap |
| Colletidae | Belowground | (Scott et al., 2011) | *Colletes* | *birkmanni* | 5 | Net |
| Colletidae | Belowground | (Scott et al., 2011) | *Colletes* | *nudus* | 1 | Net |
| Colletidae | Aboveground | (Richards et al., 2011) | *Hylaeus* | *affinis-modestus* | 1 | Net |
| Halictidae | Belowground | (Carril & Wilson, 2021) | *Agapostemon* | *angelicus-texanus* | 12 | Net, pan trap |
| Halictidae | Belowground | (Carril & Wilson, 2021) | *Agapostemon* | *obliquus-sericeus* | 9 | Pan trap, vane trap |
| Halictidae | Belowground | (Carril & Wilson, 2021) | *Agapostemon* | *splendens* | 29 | Net, pan trap, vane trap |
| Halictidae | Belowground | (Sakagami & Michener, 1962) | *Augochlorella* | *gratiosa* | 3 | Pan trap |
| Halictidae | Belowground | (Gonçalves, 2019) | *Augochlorella* | *karankawa* | 1 | Net |
| Halictidae | Belowground | (Blitzer et al., 2016) | *Augochloropsis* | *metallica* complex | 6 | Net, pan trap, vane trap |
| Halictidae | Belowground | (Carril & Wilson, 2021) | *Dieunomia* | *heteropoda* | 1 | Net |
| Halictidae | Belowground | (Wuellner, 1999) | *Dieunomia* | *triangulifera* | 1 | Pan trap |

| **Family** | **Nest Location** | **Source** | **Genus** | **Species** | **Abundance** | **Collection Method(s)** |
|---|---|---|---|---|---|---|
| Halictidae | Belowground | (Richards & Packer, 1996) | *Halictus* | *ligatus* | 38 | Net, pan trap, vane trap |
| Halictidae | Belowground | (Carril & Wilson, 2021) | *Halictus* | *parallelus* | 1 | Net |
| Halictidae | Belowground | (Cane, 2015) | *Halictus* | *rubicundus* | 1 | Net |
| Halictidae | Belowground | (Carril & Wilson, 2021) | *Lasioglossum* | *bruneri* | 1 | Vane trap |
| Halictidae | Belowground | (Carril & Wilson, 2021) | *Lasioglossum* | cf. *aliud* | 8 | Net, pan trap, vane trap |
| Halictidae | Belowground | (Michener, 2007) | *Lasioglossum* | *coactus* | 128 | Net, pan trap, vane trap |
| Halictidae | Belowground | (Carril & Wilson, 2021) | *Lasioglossum* | *connexum* | 7 | Net, pan trap, vane trap |
| Halictidae | Belowground | (Carril & Wilson, 2021) | *Lasioglossum* | *disparile* | 116 | Net, pan trap, vane trap |
| Halictidae | Belowground | (Carril & Wilson, 2021) | *Lasioglossum* | *hudsoniellum* | 26 | Net, pan trap, vane trap |
| Halictidae | Belowground | (Michener, 1974) | *Lasioglossum* | *imitatum* | 1 | Net |
| Halictidae | Belowground | (Carril & Wilson, 2021) | *Lasioglossum* | *laevissimum* | 1 | Pan trap |
| Halictidae | Belowground | (Carril & Wilson, 2021) | *Lasioglossum* | *longifrons* | 10 | Net, pan trap |
| Halictidae | Belowground | (Carril & Wilson, 2021) | *Lasioglossum* | *pilosum* | 2 | Net, vane trap |
| Halictidae | Belowground | (Carril & Wilson, 2021) | *Lasioglossum* | *pruinosum* | 2 | Net, pan trap |
| Halictidae | Belowground | (Carril & Wilson, 2021) | *Lasioglossum* | *semicaeruleum* | 42 | Net, pan trap, vane trap |
| Halictidae | Belowground | (Carril & Wilson, 2021) | *Lasioglossum* | sp24A | 2 | Pan trap |

| Family | Nest Location | Source | Genus | Species | Abundance | Collection Method(s) |
|---|---|---|---|---|---|---|
| Halictidae | Belowground | (Carril & Wilson, 2021) | *Lasioglossum* | sp2A | 4 | Pan trap |
| Halictidae | Belowground | (Carril & Wilson, 2021) | *Lasioglossum* | sp3A | 107 | Net, pan trap, vane trap |
| Halictidae | Belowground | (Carril & Wilson, 2021) | *Lasioglossum* | sp3B | 1 | Net |
| Halictidae | Belowground | (Carril & Wilson, 2021) | *Lasioglossum* | sp4A | 4 | Net, pan trap, vane trap |
| Halictidae | Belowground | (Carril & Wilson, 2021) | *Lasioglossum* | sp5A | 1 | Net |
| Halictidae | Belowground | (Carril & Wilson, 2021) | *Lasioglossum* | sp6A | 7 | Vane trap |
| Halictidae | Belowground | (Carril & Wilson, 2021) | *Lasioglossum* | spp. | 5 | Net, pan trap |
| Halictidae | Belowground | (Carril & Wilson, 2021) | *Lasioglossum* | spRed1 | 1 | Pan trap |
| Halictidae | | | | spp. | 2 | Net, pan trap, vane trap |
| Megachilidae | Aboveground | (Baker et al., 1985) | *Anthidiellum* | *notatum* | 1 | Net |
| Megachilidae | Belowground | (Yanega, 1994) | *Atoposmia* | *maryae* | 1 | Net |
| Megachilidae | Belowground | (Yanega, 1994) | *Atoposmia* | sp1A | 2 | Net |
| Megachilidae | Belowground | (Yanega, 1994) | *Atoposmia* | sp2A | 1 | Net |
| Megachilidae | Aboveground | (Fischer, 1955) | *Heriades* | *variolosa* | 3 | Net |
| Megachilidae | Aboveground | (Rozen & Hall, 2014) | *Lithurgopsis* | *apicalis* | 1 | Vane trap |
| Megachilidae | Aboveground | (B. N. Danforth et al., 2019) | *Lithurgopsis* | *littoralis* | 1 | Net |
| Megachilidae | Belowground | (Harmon-Threatt, 2020) | *Megachile* | *albitarsis* | 1 | Net |

| **Family** | **Nest Location** | **Source** | **Genus** | **Species** | **Abundance** | **Collection Method(s)** |
|---|---|---|---|---|---|---|
| Megachilidae | Belowground | (Michener, 1953) | *Megachile* | *brevis* | 8 | Net, vane trap |
| Megachilidae | Belowground | (B. N. Danforth et al., 2019) | *Megachile* | *campanulae* | 1 | Net |
| Megachilidae | Belowground | (Harmon-Threatt, 2020) | *Megachile* | *inimica* | 1 | Vane trap |
| Megachilidae | Belowground | (H. J. Williams et al., 1986) | *Megachile* | *integra* | 1 | Net |
| Megachilidae | Belowground | (B. N. Danforth et al., 2019) | *Megachile* | *montivaga* | 2 | Net, vane trap |
| Megachilidae | Belowground | (B. N. Danforth et al., 2019) | *Megachile* | *mucida* | 1 | Vane trap |
| Megachilidae | Belowground | (B. N. Danforth et al., 2019) | *Megachile* | *policaris* | 2 | Net |
| Megachilidae | Belowground | (B. N. Danforth et al., 2019) | *Megachile* | *pugnata* | 1 | Net |
| Megachilidae | Aboveground | (Pitts-Singer & Cane, 2011) | *Megachile* | *rotundata* | 7 | Net |
| Megachilidae | Aboveground | (Rozen & Hall, 2011) | *Osmia* | *chalybea* | 2 | Net |
| Megachilidae | Aboveground | (Cane et al., 2007) | *Osmia* | *subfasciata* | 1 | Net |
| Megachilidae | Aboveground | (Harmon-Threatt, 2020) | *Stelis* | *diversicolor* | 1 | Net |
| Megachilidae | Aboveground | (Michener, 1955) | *Stelis* | *lateralis* | 1 | Net |
| Melittidae | Belowground | (Carril & Wilson, 2021) | *Hesperapis* | sp1A | 1 | Net |

**Table S7:** Multivariate analysis of impacts of habitat characteristics on ground-nesting bee community composition. Results show likelihood ratio tests based on Montecarlo sampling with 999 bootstrap iterations.

| Species | Intercept $\chi^2$ | Intercept p-value | Floral abundance $\chi^2$ | Floral abundance p-value | Floral richness $\chi^2$ | Floral richness p-value | Bare ground cover $\chi^2$ | Bare ground cover p-value | Soil sand content $\chi^2$ | Soil sand content p-value | Soil depth $\chi^2$ | Soil depth p-value | Soil compaction $\chi^2$ | Soil compaction p-value |
|---|---|---|---|---|---|---|---|---|---|---|---|---|---|---|
| Overall | 512.28 | 0.25 | 37.98 | 1.00 | 94.48 | 0.98 | 58.50 | 0.98 | 58.73 | 0.95 | 62.21 | 0.25 | 54.05 | 0.28 |
| *Agapostemon angelicus-texanus* | 0.01 | 1.00 | 0.01 | 1.00 | 18.21 | 0.99 | 0.001 | 0.99 | 0.04 | 0.97 | 14.29 | 0.46 | 11.15 | 0.47 |
| *Agapostemon obliquus-sericeus* | 7.04 | 1.00 | 0.30 | 1.00 | 1.19 | 0.99 | 1.65 | 0.99 | 1.31 | 0.97 | 0.06 | 0.95 | 0.55 | 0.98 |
| *Agapostemon splendens* | 2.39 | 1.00 | 0.58 | 1.00 | 0.08 | 0.99 | 0.10 | 0.99 | 0.05 | 0.97 | 0.50 | 0.95 | 0.01 | 0.98 |
| *Anthophora phenax* | 18.90 | 1.00 | 0.00 | 1.00 | 6.41 | 0.99 | 0.88 | 0.99 | 0.00 | 0.97 | 0.02 | 0.95 | 0.00 | 1.00 |
| *Atoposmia* sp1A | 23.64 | 1.00 | 0.01 | 1.00 | 3.07 | 0.99 | 0.00 | 0.99 | 0.002 | 0.97 | 0.00 | 0.95 | 0.001 | 0.98 |
| *Augochlorella gratiosa* | 22.98 | 1.00 | 0.01 | 1.00 | 0.01 | 0.99 | 0.01 | 0.99 | 0.02 | 0.97 | 0.15 | 0.95 | 0.02 | 0.98 |
| *Augochloropsis metallica* complex | 12.02 | 1.00 | 0.21 | 1.00 | 1.16 | 0.99 | 0.04 | 0.99 | 0.45 | 0.97 | 0.42 | 0.95 | 2.11 | 0.98 |
| *Bombus griseocollis* | 21.71 | 1.00 | 0.001 | 1.00 | 0.01 | 0.99 | 0.00 | 0.99 | 2.47 | 0.97 | 0.01 | 0.95 | 0.004 | 0.98 |
| *Bombus pensylvanicus* | 21.66 | 1.00 | 0.03 | 1.00 | 1.39 | 0.99 | 4.00 | 0.99 | 5.57 | 0.97 | 0.003 | 0.95 | 0.04 | 0.98 |
| *Colletes birkmanni* | 18.85 | 1.00 | 0.00 | 1.00 | 10.23 | 0.99 | 0.02 | 0.99 | 8.52 | 0.97 | 0.01 | 0.95 | 0.02 | 0.98 |
| *Eucera speciosa* | 22.24 | 1.00 | 0.001 | 1.00 | 1.61 | 0.99 | 1.20 | 0.99 | 0.004 | 0.97 | 0.003 | 0.95 | 7.03 | 0.71 |
| *Halictus ligatus* | 0.004 | 1.00 | 2.55 | 1.00 | 0.18 | 0.99 | 0.32 | 0.99 | 0.68 | 0.97 | 0.21 | 0.95 | 1.72 | 0.98 |
| *Lasioglossum* cf. *aliud* | 18.71 | 1.00 | 7.96 | 1.00 | 11.81 | 0.99 | 8.20 | 0.99 | 0.04 | 0.97 | 0.11 | 0.95 | 0.09 | 0.98 |

| Species | Intercept $\chi^2$ | Intercept p-value | Floral abundance $\chi^2$ | Floral abundance p-value | Floral richness $\chi^2$ | Floral richness p-value | Bare ground cover $\chi^2$ | Bare ground cover p-value | Soil sand content $\chi^2$ | Soil sand content p-value | Soil depth $\chi^2$ | Soil depth p-value | Soil compaction $\chi^2$ | Soil compaction p-value |
|---|---|---|---|---|---|---|---|---|---|---|---|---|---|---|
| *Lasioglossum coactus* | 7.64 | 1.00 | 11.05 | 1.00 | 0.02 | 0.99 | 8.52 | 0.99 | 16.54 | 0.97 | 10.74 | 0.50 | 9.29 | 0.55 |
| *Lasioglossum connexum* | 6.22 | 1.00 | 2.87 | 1.00 | 0.58 | 0.99 | 0.30 | 0.99 | 2.15 | 0.97 | 2.11 | 0.95 | 1.32 | 0.98 |
| *Lasioglossum disparile* | 8.78 | 1.00 | 0.00 | 1.00 | 0.00 | 0.99 | 0.00 | 1.00 | 0.002 | 0.97 | 0.00 | 0.95 | 0.002 | 0.98 |
| *Lasioglossum hundoniellum* | 0.001 | 1.00 | 0.002 | 1.00 | 12.39 | 0.99 | 0.03 | 0.99 | 0.04 | 0.97 | 0.03 | 0.95 | 0.11 | 0.98 |
| *Lasioglossum longifrons* | 1.55 | 1.00 | 0.50 | 1.00 | 0.46 | 0.99 | 0.80 | 0.99 | 6.78 | 0.97 | 2.37 | 0.95 | 0.29 | 0.98 |
| *Lasioglossum pilosum* | 22.45 | 1.00 | 0.04 | 1.00 | 0.01 | 0.99 | 0.22 | 0.99 | 8.58 | 0.97 | 0.001 | 0.95 | 0.33 | 0.98 |
| *Lasioglossum pruinosum* | 21.17 | 1.00 | 0.002 | 1.00 | 0.01 | 0.99 | 0.001 | 0.99 | 0.01 | 0.97 | 0.04 | 0.95 | 0.004 | 0.98 |
| *Lasioglossum semicaeruleum* | 0.43 | 1.00 | 0.28 | 1.00 | 1.62 | 0.99 | 0.34 | 0.99 | 2.71 | 0.97 | 2.12 | 0.95 | 0.56 | 0.98 |
| *Lasioglossum* sp24A | 18.66 | 1.00 | 0.001 | 1.00 | 0.56 | 0.99 | 0.001 | 0.99 | 0.01 | 0.97 | 0.00 | 0.95 | 0.001 | 0.98 |
| *Lasioglossum* sp2A | 23.64 | 1.00 | 0.01 | 1.00 | 3.07 | 0.99 | 0.00 | 0.99 | 0.002 | 0.97 | 0.00 | 0.95 | 0.001 | 0.98 |
| *Lasioglossum* sp3A | 21.23 | 1.00 | 0.00 | 1.00 | 2.01 | 0.99 | 7.91 | 0.99 | 0.00 | 0.97 | 0.00 | 1.00 | 0.00 | 1.00 |
| *Lasioglossum* sp4A | 12.38 | 1.00 | 0.05 | 1.00 | 1.94 | 0.99 | 0.07 | 0.99 | 0.01 | 0.97 | 0.002 | 0.95 | 0.01 | 0.98 |
| *Lasioglossum* sp6A | 22.57 | 1.00 | 0.001 | 1.00 | 0.002 | 0.99 | 0.00 | 0.99 | 0.00 | 0.97 | 0.00 | 0.95 | 0.00 | 0.98 |
| *Megachile brevis* | 4.66 | 1.00 | 1.43 | 1.00 | 4.00 | 0.99 | 0.97 | 0.99 | 0.39 | 0.97 | 0.43 | 0.95 | 0.24 | 0.98 |
| *Megachile montivaga* | 22.39 | 1.00 | 0.02 | 1.00 | 0.04 | 0.99 | 0.001 | 0.99 | 0.01 | 0.97 | 0.003 | 0.95 | 0.01 | 0.98 |
| *Megachile policaris* | 20.62 | 1.00 | 0.00 | 1.00 | 0.001 | 0.99 | 0.001 | 0.99 | 0.01 | 0.97 | 0.002 | 0.95 | 0.01 | 0.98 |

| Species | Intercept $\chi^2$ | Intercept p-value | Floral abundance $\chi^2$ | Floral abundance p-value | Floral richness $\chi^2$ | Floral richness p-value | Bare ground cover $\chi^2$ | Bare ground cover p-value | Soil sand content $\chi^2$ | Soil sand content p-value | Soil depth $\chi^2$ | Soil depth p-value | Soil compaction $\chi^2$ | Soil compaction p-value |
|---|---|---|---|---|---|---|---|---|---|---|---|---|---|---|
| *Melissodes agilis* | 19.72 | 1.00 | 0.01 | 1.00 | 5.08 | 0.99 | 10.88 | 0.99 | 0.01 | 0.97 | 13.31 | 0.47 | 8.97 | 0.55 |
| *Melissodes boltoniae-tinctus* | 22.39 | 1.00 | 0.02 | 1.00 | 0.04 | 0.99 | 0.001 | 0.99 | 0.01 | 0.97 | 0.003 | 0.95 | 0.01 | 0.98 |
| *Melissodes communis-tepaneca* | 16.32 | 1.00 | 9.80 | 1.00 | 4.54 | 0.99 | 5.08 | 0.99 | 2.34 | 0.97 | 12.40 | 0.49 | 9.49 | 0.55 |
| *Melissodes comptoides* | 22.67 | 1.00 | 0.01 | 1.00 | 0.004 | 0.99 | 0.03 | 0.99 | 0.001 | 0.97 | 0.001 | 0.95 | 0.06 | 0.98 |
| *Melissodes coreopsis* | 8.72 | 1.00 | 0.09 | 1.00 | 0.33 | 0.99 | 6.94 | 0.99 | 0.001 | 0.97 | 2.89 | 0.95 | 0.62 | 0.98 |
| *Xenoglossa pruinosa* | 21.66 | 1.00 | 0.00 | 1.00 | 2.43 | 0.99 | 0.00 | 0.99 | 0.004 | 0.97 | 0.00 | 0.95 | 0.02 | 0.98 |

**Table S8:** Impacts of grazing rest period on habitat characteristics, assessed via likelihood ratio or F tests. For the F test (soil compaction), residual degrees of freedom are in the Intercept row. Terms significant at $\alpha < 0.05$ are highlighted in green.

| **Habitat characteristic** | **Model term** | **Coefficient** | **Coefficient Standard Error** | **$\chi^2$ or F** | **df** | **p-value** | **Fall vs. Spring p-value** | **Fall vs. Summer p-value** | **Spring vs. Summer p-value** |
|---|---|---|---|---|---|---|---|---|---|
| Floral abundance | Intercept | 6.86 | 0.41 | | | | | | |
| | Grazing rest period | 0.38 | 0.16 | 6.34 | 1 | 0.01 | | | |
| | Season - spring | -0.50 | 0.35 | 4.08 | 2 | 0.13 | 0.30 | 0.15 | 0.52 |
| | Season - summer | -0.75 | 0.38 | | | | | | |
| Floral richness | Intercept | 13.15 | 1.83 | | | | | | |
| | Grazing rest period | 2.70 | 0.68 | 14.31 | 1 | 0.0002 | | | |
| | Season - spring | 5.68 | 1.58 | 10.00 | 2 | 0.007 | 0.002 | < 0.001 | 0.19 |
| | Season - summer | 7.70 | 1.51 | | | | | | |
| Bare ground cover | Intercept | -4.62 | 0.46 | | | | | | |
| | Grazing rest period | 0.05 | 0.02 | 5.27 | 1 | 0.02 | | | |
| | Season – spring | 0.76 | 0.12 | 13.14 | 2 | 0.001 | < 0.001 | < 0.001 | 0.36 |
| | Season - summer | 0.86 | 0.12 | | | | | | |
| Soil compaction | Intercept | 1.39 | 0.10 | | 7 | | | | |

| | | | | | | |
|---|---|---|---|---|---|---|
| | Grazing rest period | 0.001 | 0.001 | 1.61 | 1 | 0.25 |

**Figure S1:** Study sites (white circles) and sampling design (inset). Sites are located in the Cross Timbers ecoregion, northwest of the Dallas-Fort Worth Metroplex. Sampling design schematic shows transects (dashed lines) and vegetation (“veg”) quadrat, soil sample, and bee trap locations. Sampling design schematic not to scale.

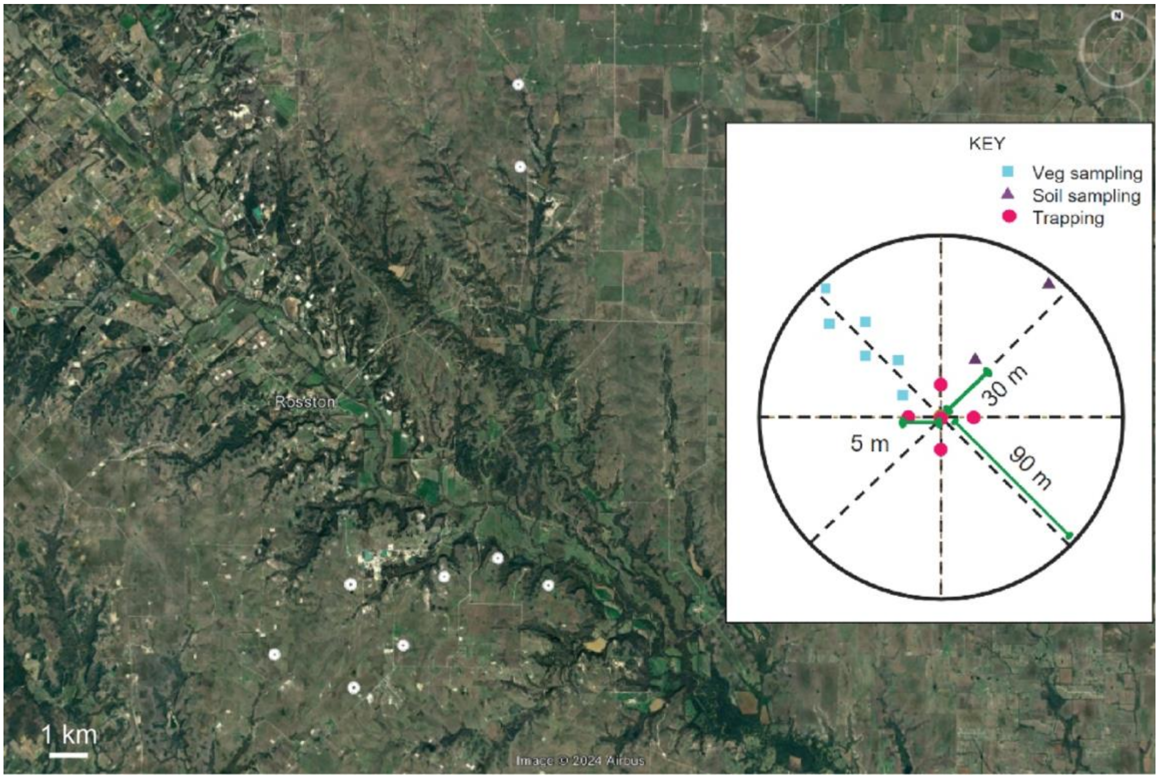

**Figure S2:** Community composition of bee assemblages at each site. Stacked bars show the proportion of bees from each genus. Genera where all species nest below ground are colored brown, aboveground-nesting genera colored green, and the genus with species that nest above and below ground with blue.

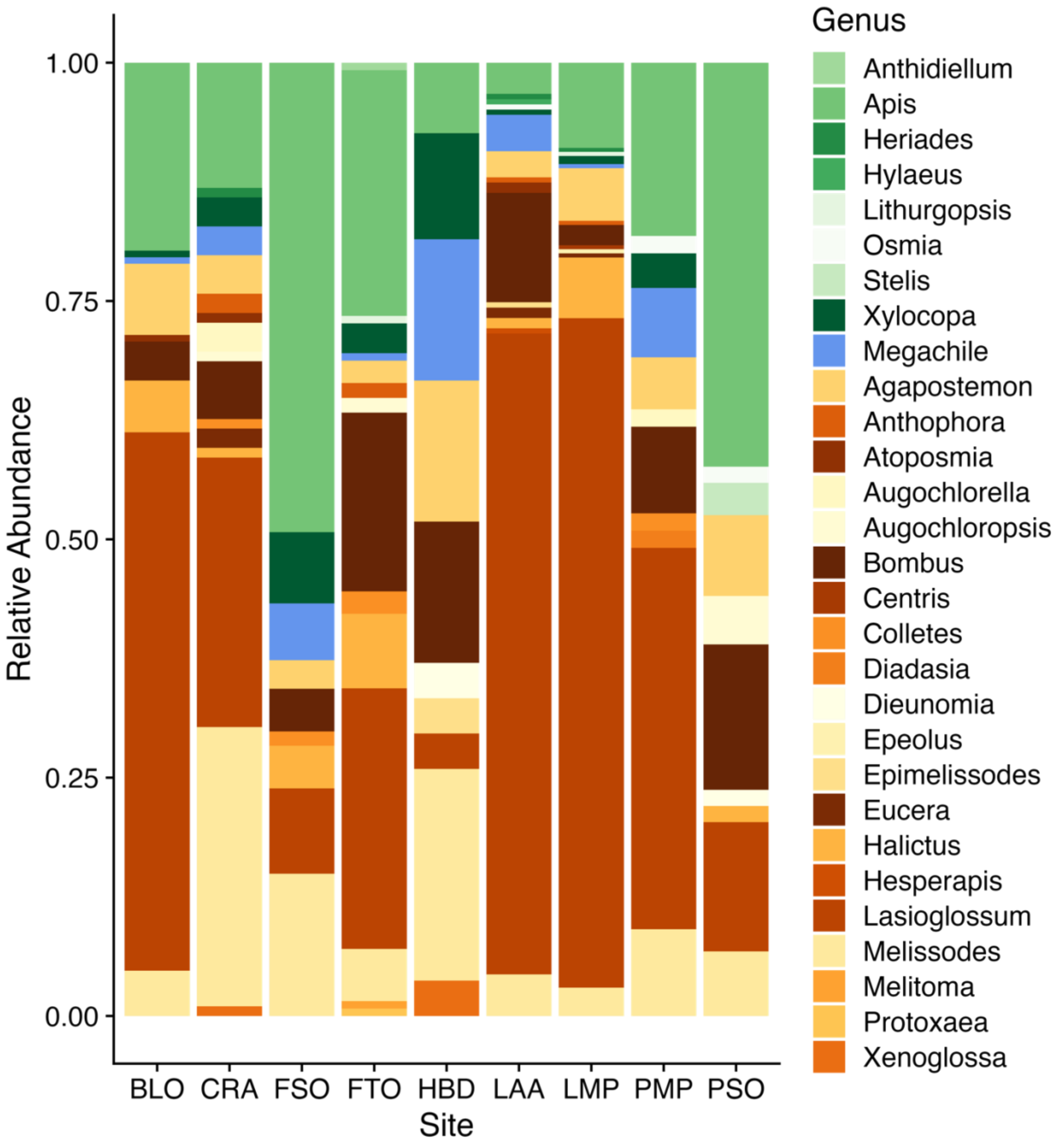
